# Chromatin Remodeling and Transcriptional Silencing Define the Dynamic Innate Immune Response of Tissue Resident Macrophages After Burn Injury

**DOI:** 10.1101/2025.05.15.654340

**Authors:** Han G. Kim, Marie-Pierre L. Gauthier, Aidan Higgs, Denise A. Hernandez, Mingqi Zhou, Jason O. Brant, Rhonda L. Bacher, Dijoia B. Darden, Shannon M. Wallet, Clayton E. Mathews, Philip A. Efron, Michael P. Kladde, Robert Maile

**Affiliations:** Sepsis and Critical Illness Research Center, Department of Surgery, College of Medicine, University of Florida, Gainesville, Florida, United States; Department of Biochemistry and Molecular Biology, College of Medicine, University of Florida, Gainesville, Florida, United States; National Institute of Mental Health Intermural Research Program, National Institutes of Health, Bethesda, Maryland United States; Department of Biostatistics, College of Public Health & Health Professions, University of Florida, Gainesville, Florida, United States; University of Florida Health Cancer Center, University of Florida, Gainesville, Florida, United States; Department of Oral Biology, College of Dentistry, University of Florida, Gainesville, Florida, United States; Department of Pathology, Immunology and Laboratory Medicine, College of Medicine, University of Florida, Gainesville, Florida, USA; Department of Infectious Diseases and Immunology, College of Veterinary Medicine, University of Florida, Gainesville, Florida, USA

## Abstract

Severe burn injury induces long-lasting immune dysfunction, but the molecular mechanisms underlying this phenomenon remain unclear. We hypothesized that burn injury leads to epigenetic and transcriptional reprogramming of innate immune cells. Splenic F4/80⁺ macrophages were isolated from mice at days 2, 9, and 14 days post-20% contact burn injury. Targeted transcriptomics and MAPit single-molecule chromatin profiling were used to assess immune, metabolic, and epigenetic changes. Canonical pathway analysis was performed to infer functional shifts over time. Burn injury induced a biphasic response in macrophages. Early after injury (Day 2), there was broad transcriptional suppression and epigenetic silencing of inflammatory regulators, including *Stat3*, *Traf6*, and *Nfkb1*. Over time (Days 9 and 14), loci associated with anti-inflammatory mediators such as *Il-10* and *Socs3* exhibited progressive chromatin opening and transcriptional upregulation. Metabolic gene profiles revealed persistent suppression of mitochondrial and oxidative phosphorylation programs. Canonical pathway analysis demonstrated early IL-10 signaling activation with sustained suppression of classical macrophage activation pathways. Chromatin architecture changes included nucleosome sliding and ejection events, consistent with dynamic, locus-specific regulation. This work challenges the classical notion of burn-induced immune suppression as purely a consequence of systemic inflammation. Instead, we reveal a programmed and locus-specific epigenetic architecture that may shape macrophage immune and metabolic function long after the acute phase.

## Introduction

Severe burn injury triggers a complex immune response that can lead to life-threatening infections and sepsis. According to the American Burn Association, approximately 450,000 patients receive treatment for burns each year, with mortality rates of major burns being around 22.5%^1^. The prevailing model proposes an initial exaggerated inflammatory cytokine storm which appears to be driven by Damage-Associated Molecular Patterns (DAMPs), such as hemagglutinin (HA) and circulating double-stranded (ds) DNA after rapid glucocorticoid-mediated systemic death of immune cells. Patients who survive this phase often exhibit an immunological endotype of Persistent Inflammation, Immunosuppression, and Catabolism Syndrome (PICS), which clinically presents as chronic immunosuppression, low-grade inflammation, and lean tissue wasting^2–11^. It is important to note that rather than sequential discrete immune response phenotypes, pro- and anti-inflammatory mediators are detected simultaneously in <u>both</u> of these clinical phases^2, 12–16^, often described as a mixed antagonist response syndrome (MARS^12, 17–22^). Many groups have attempted to delineate the underlying mechanisms in order to predict and prevent onset of PICS.

The emerging concept of innate immune memory encompasses two opposing phenotypes: trained immunity versus immune tolerance^23^. Trained immunity is characterized by a heightened inflammatory response upon rechallenge, due to sustained priming of innate cells. In contrast, tolerance is a state of refractoriness, with blunted pro-inflammatory cytokine production and a bias toward anti-inflammatory mediators (like IL-10). Trained immunity and tolerance are both driven by long-term reprogramming of innate immune cells through <u>metabolic</u> and <u>epigenetic</u> mechanisms. Despite comprehensive epigenomic analysis in sepsis models^24^ and subsequent linkage with the resultant cellular inflammatory response, there remains a paucity of comparable research in burn.

In this Brief Report, we demonstrate that specific time-dependent alterations in the immune and metabolic transcriptomic responses of F4/80+ tissue-resident macrophages (trMø) correspond to significant time-dependent epigenetic changes in a mouse model of severe cutaneous burn injury.

## Methods

### Mouse cutaneous burn injury

Female C57BL/6 mice weighing 18-22 g (8-12 weeks old) were used for all experiments. Animals were anesthetized by an intraperitoneal (*i.p.*) injection of 0.024 mL/g of Avertin and had their dorsal and flank hair clipped. A subcutaneous injection of Ethiqa XR® (buprenorphine extended-release injectable suspension, 1.3 mg/ml; Fidelis Animal Health, Inc., NJ, USA) was administered prior to injury for pain control. To generate a full-thickness thermal burn of approximately 20% of the Total Body Surface Area (TBSA), a 65 g copper rod (1.9 cm in diameter) was heated to 100°C and applied to four separate areas, each for 10 sec, to the animal’s dorsum and flank at a standard pressure that we have demonstrated produces a full-thickness burn ^25^. The mice were allowed to recover on a heating pad and were resuscitated with an *i.p.* injection of lactated Ringer’s solution (2.5 mL). Once resuscitated, the mice were placed in individual cages, provided food *ad libitum* and maintained on Ethiqa XR®, and continuously monitored until euthanasia. Sham controls underwent all procedures in parallel, except for the burn injury. Mice were euthanized 2, 9, or 14 days after sham or burn injury. All animal work was performed under University of Florida Institutional Animal Care and Use Committee (IACUC)-approved protocols.

### F4/80+ tissue-resident macrophage (trMφ) purification

Spleens were removed from the euthanized mice, and following mechanical dissociation and standard Ammonium-Chloride-Potassium (ACK) red blood cell lysis, single-cell suspensions were filtered through a 70 µm mesh and centrifuged at 300 *× g* for 10 min. Cells were resuspended in Magnetic-Activated Cell Sorting (MACS) buffer (phosphate-buffer saline (PBS), pH 7.2, containing 0.5% bovine serum albumin (BSA) and 2 mM ethylenediaminetetraacetic acid (EDTA), and total cell counts were determined using a hemocytometer with trypan blue exclusion. Fifty million splenocytes were enriched for F4/80^+^ using mouse F4/80 magnetic bead (130-110-443, Miltenyi Biotec, Teterow, Germany) selection through LS columns (130-091-596, Miltenyi Biotec) on a QuadroMACS setup (Miltenyi Biotec), according to manufacturer’s instructions. One spleen routinely yielded approximately 2 x 10^6^ cells with >95% purity ^25, 26^. Enriched F4/80^+^ cells were used immediately for transcriptomic and epigenetic analysis.

### RNA Extraction and Transcriptomic Analysis

Isolation of mRNA was performed using mRNA easy kits according to manufacturer’s instructions. mRNA was quantified using a Nanodrop 2000^TM^ spectrophotometer (Waltham, MA). NanoString technology and the nCounter Mouse Immunology Panel (XT-CSO-MIM1-12, NanoString, Seattle, WA) and Mouse Metabolic Panel (XT-CSO-MMP1-12) were used to simultaneously evaluate 561 immune- and 768 metabolic-related mRNAs in each sample, respectively ^27^. Each sample was analyzed in triplicate according to manufacturer’s instructions. nSolver v4.0 and ROSALIND integrated analysis platforms were used to generate appropriate data normalization as well as fold-changes, resulting ratios, and differential expression. Ingenuity Pathway Analysis (IPA: Qiagen, USA) and R statistics were used to identify canonical pathway-specific responses^27^.

### Methyltransferase Accessibility Protocol for Individual Templates combined with Flap-Enabled Next-Generation Capture (MAPit-FENGC) epigenetic analysis of burn injury-induced trMø

MAPit-FENGC is a validated method that allows for multiplexed, simultaneous single-molecular level analysis of methylation and accessibility at target promoter and enhancer regions^28^. The full FENGC methodology can be found at Zhou et al. (2022)^28^and detailed in **Supplemental Materials 1.**

Target gene promoters and enhancers of interest were located using transcription start site (TSS) region and ENCODE candidate of *cis*-regulatory element annotations, respectively, from the mm10 genome assembly. Flap oligos 1 and 2 as well as nested oligos 3 are catalogued in **Supplementary Materials 2**. All three oligos (4 nmole each) were ordered as 0.3-mL each as salt-free in 96-well format (Eurofins Genomics, KY, USA). Purified amplicons were submitted to the University of Florida Interdisciplinary Center for Biotechnology Research (UF-ICBR, RRID:SCR_019152) for barcoding and SMRT bell library construction. Libraries were sequenced on a PacBio SEQUEL IIe (Menlo Park, CA) instrument by UF-ICBR. The library pool was loaded at 120 pM, using diffusion loading and 20- to 30-h movies with HiFi generation and demultiplexing. Sequencing Kit 2.0 (PacBio, 101-389-001) and Instrument Chemistry Bundle Version 11.0 were used. All other steps were performed using recommended protocol by the PacBio sequencing calculator. For epigenetic analysis, high-fidelity circular consensus sequencing (HiFi CCS) was generated using default parameters. CCS reads were filtered for ≥5 single polymerase read passes and aligned to the reference sequences using the python reAminator pipeline^29^. Cut-offs of ≥ 95% conversion rate and alignment of ≥ 95% length of each reference sequence were applied. To distinguish endogenous CG methylation and M.CviPI-probed GC methylation unambiguously, GCG sites were removed, resulting in calculation and plotting of HCG and GCH methylation (where H is A, C, or T). Averaged heatmaps were constructed based on averaged percentages HCG/GCH per nucleotide post-methylscaper analysis^30^.

A total of 588 primers for a total of 196 selected promoter and enhancer regions of immune and trauma-related genes (“BurnMAP”; **Supplemental Materials 2**) were used to concurrently profile chromatin accessibility and methylation. Regions with less than three amplified samples in sham, D2, or D9 or <100 reads per condition were not considered. TSS location was determined by RefSeq annotations for each gene. For the multiplexed analysis, regions were trimmed to 100 bp upstream of the TSS site. The average accessibility of each sequence 100 bp upstream of the TSS was then calculated for each read. The average accessibility was then taken for each read. Genes with low GCH resolution around the TSS site and genes that did not pass Levene’s test for equality of variances were cut. Combined with QC filtering, this resulted in 98 final gene loci. A *t*-test was conducted for p-values as well as log_2_ fold-changes of accessibility log_2_ fold-changes of genes that had significant p-values (< 0.05) were graphed using PRISM (GraphPad Software, Inc v10.1). A two-way ANOVA with Geisser-Greenhouse correction and *post-hoc* Tukey’s was conducted for the clustering analysis between the sham, D2, and D9 mice.

## Results and Discussion

### Transcriptomic Reprogramming of Splenic trMø Following Burn Injury

Wild-type C57BL/6 mice underwent a 20% TBSA full-thickness cutaneous contact burn or sham injury (n = 6 per group). F4/80^+^ enriched cells were purified from spleens collected at acute (Day 2, D2), sub-chronic (Day 9, D9), and chronic (Day 14, D14) time points after burn injury. RNA was subsequently purified and used to perform immune and metabolic gene transcriptomic analysis with corresponding Ingenuity Pathway Analysis (IPA). **Figure 2** illustrates the immune and metabolic (**Fig 1A**) gene expression changes in the burn groups compared to compared to sham injured mice. Enriched F4/80^+^ cells at all time points after burn injury exhibit significant (p < 0.05) up- and down-regulation of a broad range of immune and metabolic genes (DEG) compared to uninjured sham mice (the top 50 significantly altered DEG are shown in **Supplemental Table 1**). We then analyzed the complete set of all significant (p < 0.05) metabolic and immune DEG by IPA to interrogate canonical signaling pathways at these time points after burn injury compared to sham-injured mice (**Fig 1B**). Multiple canonical signaling pathways were significantly impacted. For example, significant <u>increases</u> in IL-10 Signaling, PPAR Signaling, PTEN Signaling, MSP-RON Macrophage Signaling pathways, glycolysis (mainly via increased expression of hexokinases and glucose-6-phosphatase genes), and OXPHOS pathway Z-scores were seen at burn D2 and D14 versus sham-injured mice. Notable <u>decreases</u> in expression were seen in S100 Family Signaling, IL-6 Signaling, and Macrophage Classical Activation Signaling pathways at D2. D14 post-burn expression exhibited significant decreases in Macrophage Classical Activation Signaling Interleukin-1 Family Signaling, and INOS Signaling amongst others. At burn D9, the pathways significantly affected were less numerous, and centered mainly on upregulation of Cell Cycle and Cell Cycle Regulation-related pathways. Direct comparisons between D9 and D14 with D2 are shown in **Supplemental Figure 1.** Taken together, these transcriptomic data suggest a biphasic innate immune response in F4/80^+^-enriched splenocytes after burn injury.

**Figure 1:**
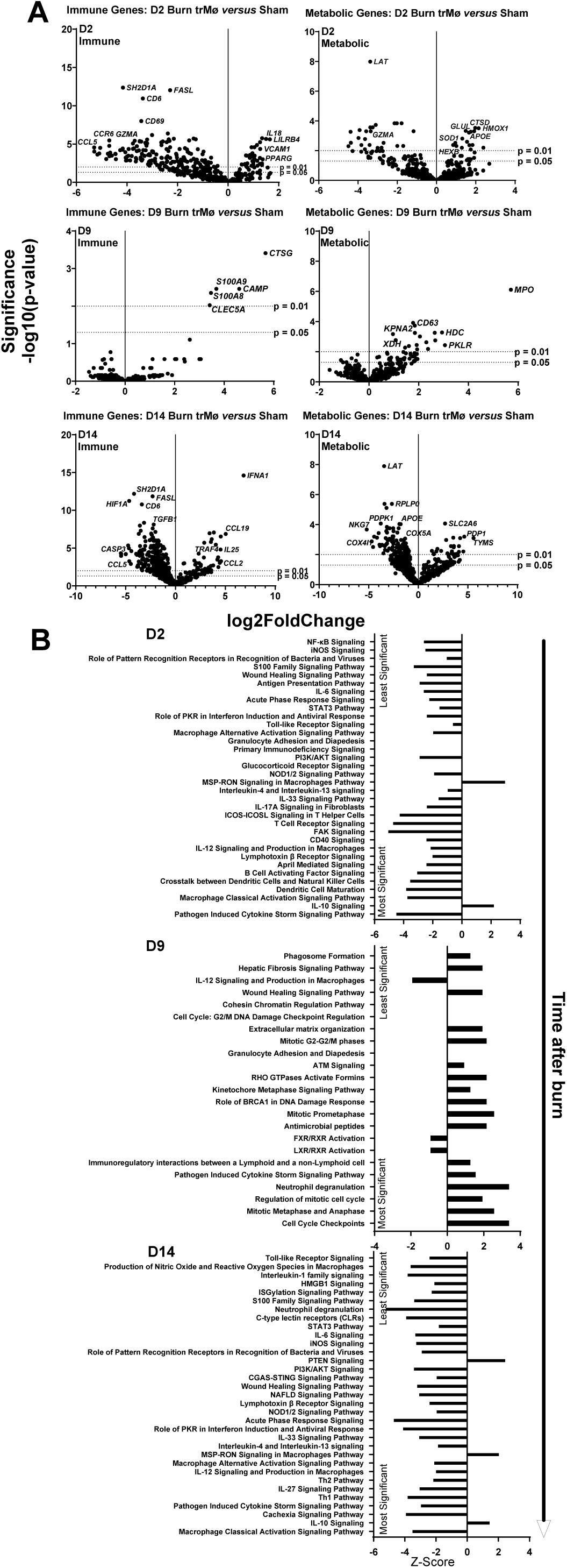
Splenic tissue-resident macrophages (trMø) harvested after burn injury exhibit specific immune and metabolic gene and pathway changes compared to trMø from sham-injured mice: mRNA isolated from splenic trMø 2, 9, or 14 days after 20% Total Body Surface Area burn injury, were tested for expression of immune and metabolic genes by nanoString analysis compared to trMø harvested from sham-injured mice. **A)** Data are presented as the log_2_-transformed differential fold change in immune or metabolic gene expression as shown in each label, with associated p-value significance (using Welch’s T test); **B)** from these data, canonical immune and metabolism pathways with associated Z-scores and significance (using Welch’s T test) were derived by Ingenuity Pathway Analysis (IPA).

**Figure 2:**
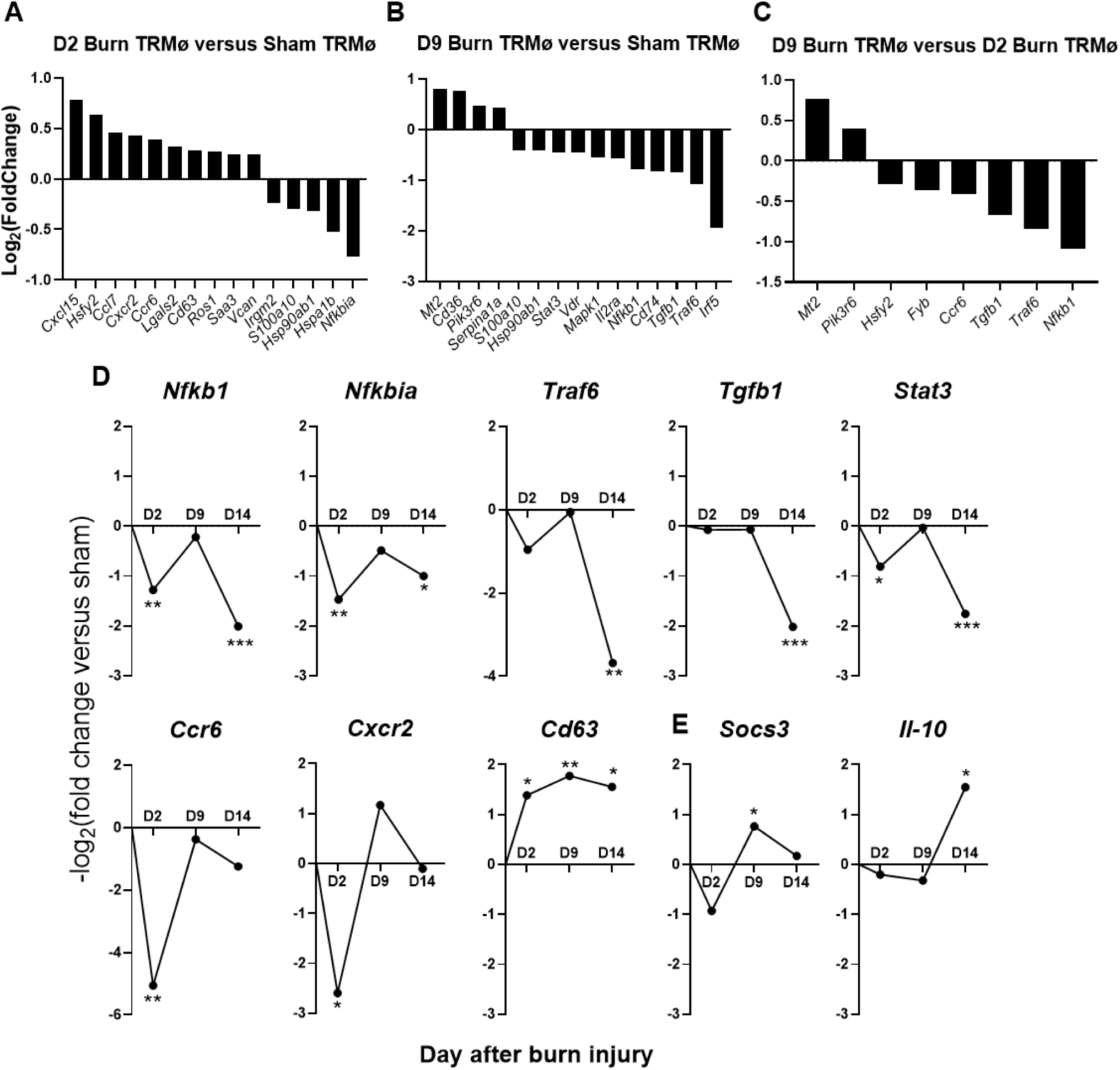
Chromatin accessibility at selected promoter and enhancer regions of immune-related genes in splenic tissue resident macrophages (TRMø) following burn injury, which is associated with significant transcriptomic changes. **(A-C)** Flap-Enabled Next-Generation Capture (FENGC) of target sequences from F4/80^+^-enriched splenocytes at Day 2 (D2) and Day 9 (D9) was performed following 20% TBSA cutaneous burn injury in C57BL/6 mice. Gene regions were selected based on TSS proximity and mm10 genome annotations. Data shown are log_2_ fold changes in chromatin accessibility from **A)** D2 burn mice versus sham-injured control, **B)** D9 burn mice versus sham, and **C)** D9 burn mice versus D2 burn mice. **(D-E)** Matching transcriptomic data for selected genes which significant log_₂_ fold changes in chromatin accessibility. mRNA isolated from splenic TRMø harvested from sham injured mice, 2, 9, or 14 days after 20% TBSA burn injury, were tested for expression of immune and metabolic genes by nanoString analysis. Data are presented as the log2-transformed differential fold change versus sham, with associated p-value significance (using Welch’s t test; *p < 0.05, **p < 0.01, ***p < 0.005).

### Epigenetic Reprogramming of Immune and Metabolic Genes in trMø Following Burn Injury

We therefore hypothesized that epigenetic changes would occur in key immune and metabolic genes in trMø at D2 and D9 post-burn compared to sham-injured mice. In parallel to the transcriptomic studies, we also purified F4/80^+^-enriched splenocyte nuclei from the same wild-type C57BL/6 mice that underwent the sham or burn injury at D2 and D9 after injury. Following isolation of nuclei, we applied MAPit single-molecule footprinting followed by FENGC target enrichment to concurrently profile chromatin accessibility and methylation over 180 selected promoter TSS and enhancer regions of the BurnMAP library. Epigenetic profiles were compared between burn and sham conditions at D2 (acute) and D9 (sub-chronic) post-injury. At D2 post-burn, significant increases in promoter/enhancer accessibility for genes such as *Cxcl15*, *Hsfy2*, *Ccl7*, and *Cxcr2* (**Fig 2A**) were observed, suggesting an early proinflammatory transcriptional poising, particularly for chemokines and stress-associated genes. In contrast, a subset of genes, including *Hspa1b*, *Hsp90ab1*, and *Nfkbia* showed reduced accessibility, potentially reflecting early repression of heat shock proteins and NF-κB regulatory feedback elements (as previously described in burn patient studies^31^). By D9 post-burn (**Fig 2B**), promoters of *Mt2*, *Cd36* and*Pik3r6* displayed significantly increased accessibility, consistent with a transition toward stress adaptation and metabolic or anti-inflammatory signaling. Conversely, regulatory regions associated with *Nfkb1*, *Tgfb1*, *Traf6*, *Stat3* and *Irf5* showed significant reductions in accessibility, indicating persistent epigenetic silencing of key inflammatory and interferon response pathways. Comparison between D9 and D2 trMø (**Fig 2C**) revealed a significant time-dependent decline in accessibility at promoters of *Nfkb1*, *Traf6*, *Tgfb1*, *Ccr6*, and *Fyb* pointing to progressive epigenetic repression of inflammatory signaling and chemotaxis-related genes. Only *Mt2* and *Pik3r6* exhibited increased accessibility over time, implicating their potential role in sustained metabolic or regulatory programs in chronically reprogrammed macrophages.

Significant log₂-fold changes in chromatin accessibility were examined in genes whose transcription was affected by burn injury (**Fig 2D**). Genes with reduced chromatin accessibility at D2 and D9, such as *Nfkbia, Nfkb1*, *Traf6*, *Stat3*, and *Tgfb1*, exhibited sustained downregulation of expression by D14, suggesting that early epigenetic silencing led to persistent transcriptional repression. We also observed transcriptional upregulation of *Socs3* and *Il-10* following burn injury (**Fig 2E**) which may reflect a shift toward an immunosuppressive macrophage phenotype, where *Socs3* inhibits pro-inflammatory signaling pathways and *Il-10* promotes sustained anti-inflammatory responses. We and others have shown that alteration in these genes to be biologically associated with the dysfunctional immune response after burn injury^32–35^

### Epigenetic Transcriptional Silencing of *Stat3* Following Burn Injury

To determine whether burn injury alters chromatin structure at key immune regulatory loci, we performed high-resolution accessibility profiling of various promoters from splenic F4/80^+^-enriched splenocytes using GCH methylation-based MAPit analysis. GCH methylation levels were used as a proxy for chromatin openness, with increased methylation indicating accessibility in nucleosome-free regions. **Figure 3A** shows that at baseline, the *Stat3* promoter exhibited moderate to high accessibility across a ∼500 bp region surrounding the TSS, with background levels of endogenous HCG methylation, consistent with an open, epigenetically permissive state. In contrast, F4/80^+^-enriched splenocytes from mice at D2 post-burn showed a clear reduction in accessibility upstream and downstream of the TSS. By D9 post-burn, accessibility was further diminished, with the lowest GCH methylation levels observed across the entire promoter region. These data indicate that *Stat3* promoter accessibility is progressively reduced following burn injury, with a temporal decline in chromatin openness from day 2 to 9. Given *Stat3*’s central role in cytokine signaling, macrophage polarization, and immune homeostasis, these findings suggest that burn-induced epigenetic silencing at the *Stat3* locus may contribute to persistent transcriptional suppression in trMø during the subacute and chronic post-injury phases (**Fig 2D**).

**Figure 3:**
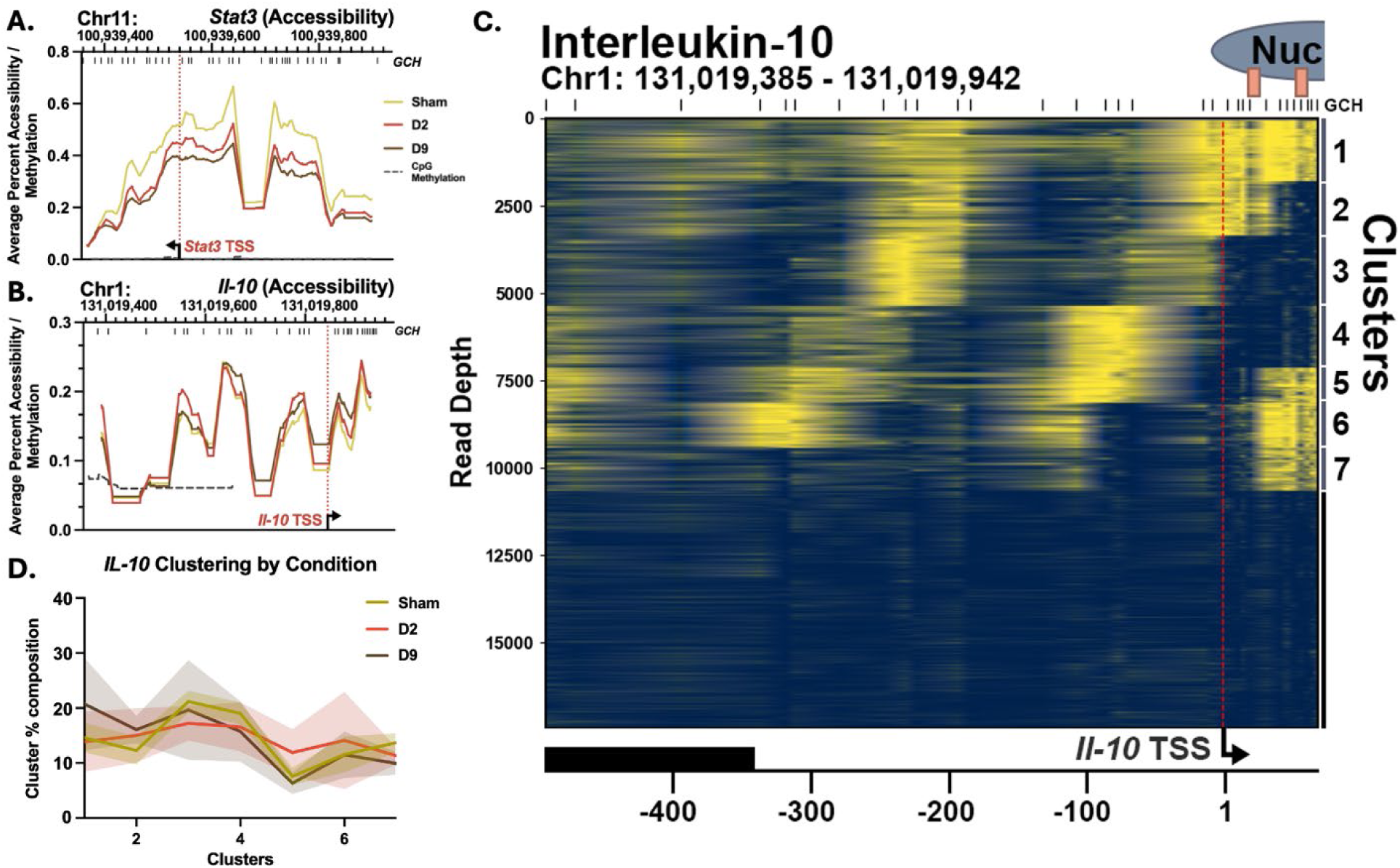
Average chromatin accessibility of the *Stat3* and *Il-10* promoter regions as well as single molecular clustering analysis of *Il-10*: Accessibility line graphs based on average GCH methylation are provided (n = 5). Groups based on days after burn are colored per figure key. Available HCG methylation data of all 15 mice is plotted by a dashed black line. Genomic coordinates and positions of GCH sites provided by vertical lines at the top of each graph. TSS position and transcript orientation is provided by the vertical dashed red line and arrow. Averaged data is presented for the **A)** *Stat3* promoter, and **B)** *Il-10* promoter. Single molecule clustering for **C)** *Il-10* is provided with clustered group labels on the right. A 147bp nucleosome reference is provided at the bottom of the graph along with relative position from the TSS. **D)** Average proportion of reads in a given cluster based on burn condition is provided for the *Il-10* promoter. Standard deviation is provided by the shaded error bands.

### Burn-Induced Chromatin Reorganization at the *Il-10* Locus Reflects a Shift Toward **Macrophage Tolerance**

Accessibility levels across the *Il-10* promoter region remain relatively consistent among all different conditions (**Fig 3B**). Using a single-molecule clustering method, we pooled all *Il-10* reads across all samples to visualize dynamic nucleosome positions across the *Il-10* promoter (**Fig 3C-D**). Molecules in clusters 7-4 show nucleosome occupancy of the *Il-10* TSS, consistent with promoter repression (**Fig 3C**). Clusters 4-1, meanwhile, show incremental, downstream nucleosome sliding, first localizing the TSS to the edge of a nucleosome (cluster 3), and sequentially exposing two actively bound transcription factor binding sites (clusters 2 and 1). A significantly lower proportion of reads (p < 0.05) in the chronic condition populated the transcriptionally inactive cluster 7 (**Supplemental Figure 2**). Furthermore, a trend towards increased average percentages of clusters 1 and 2 in the burned mice at D9 suggests that a higher proportion of cells could be in a transcription-ready epigenetic conformation. We posit that these shifts in nucleosome arrangements may indicate precursor epigenetic reprogramming that permits the robust transcriptional activity observed in D14 mice (**Fig 2E**). Finally, the D2 condition presents an even distribution of nucleosome states indicating heterogeneity of nucleosome remodeling events in earlier time points. The induction of downstream nucleosome sliding and transcription-ready chromatin at the *Il-10* promoter in F4/80^+^-enriched splenocytes by D9 post-burn suggests an epigenetically driven shift toward an anti-inflammatory state, contributing to macrophage-mediated immune suppression and chronic dysfunction after severe injury.

#### Nucleosome Dynamics Define Epigenetic Silencing at *Il2ra*, *Irak4*, *Tgfb1*, and the *Socs3* Enhancer

In addition to *Stat3* and *Il-10*, averaged accessibility plots for *Il2ra*, *Irak4*, *Tgfb1*, and a *Socs3 cis*-regulatory enhancer are presented (**Fig 4A-D**). *Il2ra* presents decreased averaged accessibility for the D9 condition around the TSS site. However, accessibility remains consistent in an upstream region of the promoter containing elevated levels of HCG methylation. *Irak4*, which plays a role in TLR4 signaling, presents decreased promoter accessibility in both D2 and D9 conditions. *Tgfb1* has decreased average accessibility for the D9 condition around the TSS site. Finally, the *Socs3* enhancer region maintains relatively consistent accessibility upstream the enhancer site but decreased D9 accessibility within the enhancer region.

**Figure 4:**
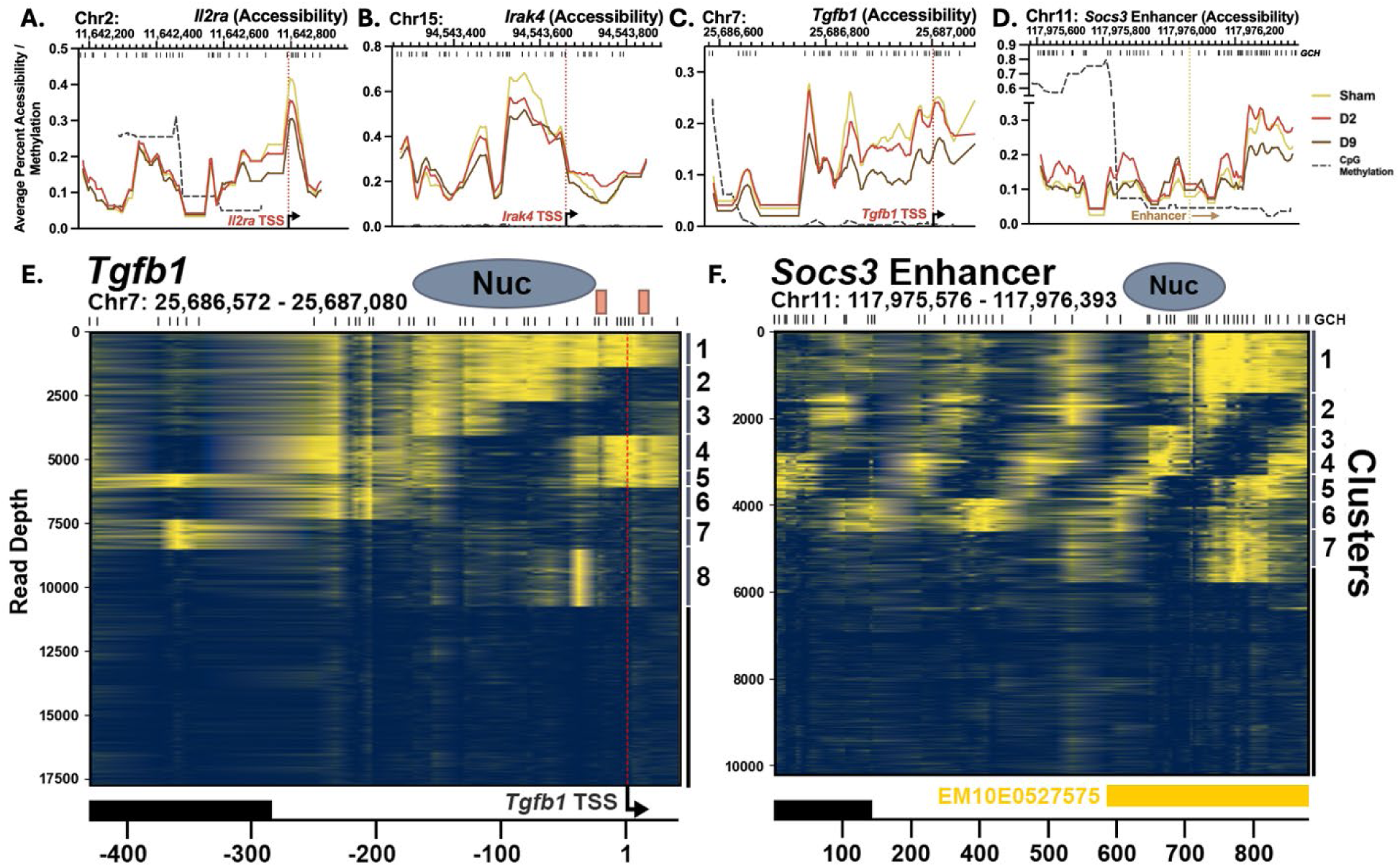
Average chromatin accessibility of *Il2ra*, *Irak4,* and *Tgfb1* promoter regions, *Socs3* enhancer regions, as well as single-molecule clustering of *Tgfb1* and the *Socs3* enhancer: Accessibility line graphs based on average GCH methylation are provided (n = 5). Groups based on days after burn are colored per figure key. Available HCG methylation data of all 15 mice is plotted by a dashed black line. Genomic coordinates and positions of GCH sites provided by vertical lines at the top of each graph. TSS position and transcript orientation is provided by the vertical dashed red line and arrow. Averaged data is presented for the **A)** *Il2ra* promoter, **B)** *Irak4* promoter, **C)** *Tgfb1* promoter, and **D)** *Socs3* enhancer. Single molecule clustering for **E)** *Tgfb1*, and **F)** the *Socs3* enhancer is provided with clustered group labels on the right. A 147bp nucleosome reference is provided at the bottom of the graph along with relative position from the TSS.

Clustering of *Tgfb1* was of note as clusters 1-4 depict a nucleosome sliding event that regulates two transcription binding sites (**Fig 4E**). Stepwise loss of *Tgfb1* promoter openness and transcription factor binding by D9 post-burn is consistent with the observed decrease in *Tgfb1* transcript at D14 (**Fig 2D**), given that changes in promoter chromatin architecture precede changes in accumulated transcript. The proportion of reads in a given cluster by condition is provided for *Tgfb1* in **Supplemental Figure 3.** Of note, the most accessible cluster 1 is significantly downregulated (p < 0.05) in the D9 condition compared to sham while a less accessible cluster 6 is significantly upregulated (p < 0.05) in the D9 condition compared to D2.

Our single-molecular analysis predicts transcriptional upregulation of *Il-10* and transcriptional downregulation of *Tgfb1* in D9 mice. Although the transcript changes of *Il-10* and *Tgfb1* are negligible at D9 (**Fig 2D-E**), we observe significant changes by D14. Therefore, our epigenetic analysis suggests early epigenetic changes that can predict transcriptomic outcomes at later time points.

Our single-molecule clustering approach was also used to visualize nucleosome conformation changes in a *cis*-regulatory enhancer region. Average HCG levels indicate high CpG methylation at the 5’ end of the selected region (**Fig 4D**). Indeed, single-molecular analysis reveals a randomly positioned array of nucleosomes with accessible linkers upstream of the enhancer (**Fig 4F**). Meanwhile, the *cis*-regulatory enhancer for *Socs3* (EM10E0527575) presents a nucleosome sliding event in the 3’ direction between clusters 7-2 followed by nucleosome ejection in cluster 1. This sliding event regulates a high-occupancy transcription factor binding site which could play a role in *Socs3* transcription (**Fig 2D**). However, there were no significant changes in the proportion of reads from sham, D2 and D9 as presented in **Supplemental Figure 4**.

## Conclusion

This study provides the first high-resolution, time-resolved integration of transcriptomic and epigenetic profiling of tissue-resident F4/80⁺ macrophages after burn injury, revealing a novel paradigm of innate immune reprogramming. Using MAPit-FENGC single-molecule accessibility mapping alongside nanoString-based transcriptional analysis, we uncover temporally coordinated chromatin remodeling events that likely drive gene expression patterns associated with immune memory. We identify epigenetic silencing of *Stat3*, *Tgfb1*, and *Nfkb1*, as well as transcriptional and structural poising of *Il-10* and *Socs3*, supporting a shift toward an anti-inflammatory, tolerogenic macrophage phenotype. These features emerge progressively post-burn, implicating early chromatin remodeling as a key determinant of immune dysfunction after burn injury. Our findings challenge the classical notion of burn-induced immune suppression as purely a consequence of systemic inflammation or central / peripheral myeloid turnover. Instead, we reveal a programmed and locus-specific epigenetic architecture that may shape macrophage immune and metabolic function long after the acute phase. This work expands current models of innate immune memory by demonstrating that trauma can also drive functional epigenetic imprints in macrophages.

Epigenetic immune memory (tolerance or training) means that the immune system does not simply reset to baseline after the acute inflammatory phase of burn. Burn and trauma patients often enter a state of chronic inflammation and immune suppression in which they are less capable of preventing infections or healing wounds. Epigenetic silencing of key inflammatory genes (tolerance) provides a mechanistic basis for this devastating clinical syndrome. On the other hand, there is evidence, including our work^32, 35–38^, that elements of trained immunity can also emerge after injury. Thus, both the tolerant and trained outcomes of innate memory are likely relevant to burns and trauma, with an initial injury simultaneously dampening certain immune functions while amplifying others *via* epigenetic changes. This may underlie both the paradox of burn and trauma survivors who suffer both recurrent infections and persistent inflammatory complications.

Several limitations of this study should be acknowledged. Although the significant differences observed were consistent across mice and supported by both chromatin accessibility and transcriptional data, larger cohorts would strengthen confidence in these conclusions and enable more granular analyses of variability among individual animals. This study focused exclusively on splenic F4/80⁺ tissue-resident macrophages, a critical but single innate immune population. It remains unknown whether similar epigenetic remodeling occurs across other innate cell subsets in this and other locations (e.g., lung, bone marrow, or peripheral) after burn injury. Integrating additional assays (e.g., CUT&RUN for histone marks, ATAC-seq, or RNA-seq) could provide a more comprehensive view of the chromatin landscape changes after trauma.

Despite these limitations, the study provides strong mechanistic evidence that burn injury drives coordinated and locus-specific epigenetic reprogramming, laying important groundwork for future larger-scale, multi-tissue, and longitudinal investigations. These new insights have broad implications not only for burn/trauma care but also for understanding immune paralysis in other trauma contexts, and may offer new targets for immunomodulatory therapies, such as interventions that re-calibrate epigenetic programming.

## Acknowledgements

We would like to thank the following sources of funding: NIH NIGMS R01GM146134 (RM/SMW).

## Supplemental

**Supplemental Figure 1:**
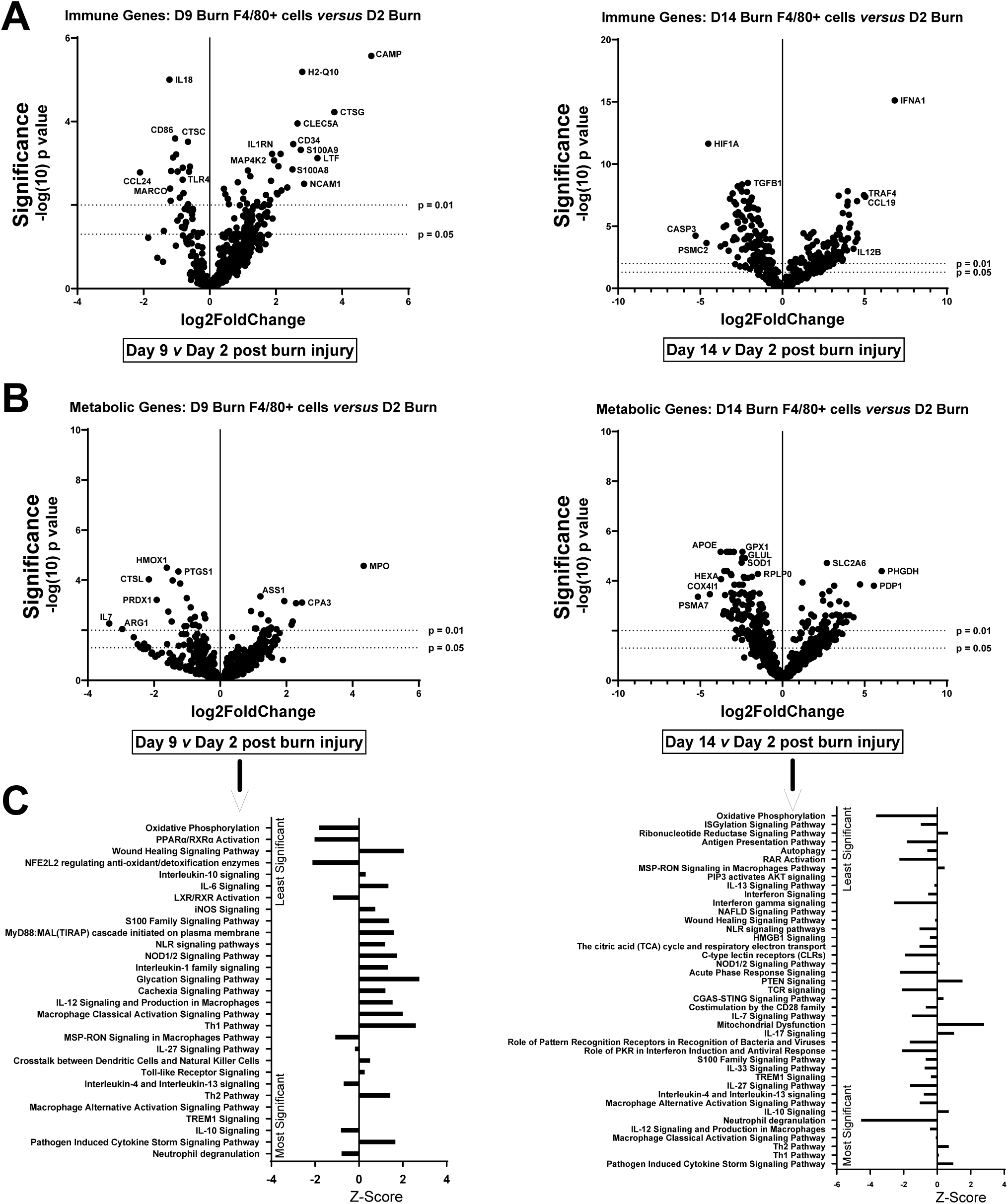
Splenic tissue resident macrophages (trMø) harvested at different time points after burn injury exhibit differential specific immune and metabolic gene, and canonical pathway changes: mRNA isolated from splenic trMø harvested 2, 9 or 14 days after 20% Total Body Surface Area burn injury, were tested for expression of immune and metabolic genes by nanoString analysis. **A-B)** Data are presented as the log2-transformed differential fold change in **(A)** immune or **(B)** metabolic gene expression as shown in each label, with associated p-value significance (using Welch’s T test); **C)** from these data, canonical immune and metabolism pathways with associated Z-scores and significance (using Welch’s T test) were derived by Ingenuity Pathway Analysis (IPA).

**Supplemental Figure 2:**
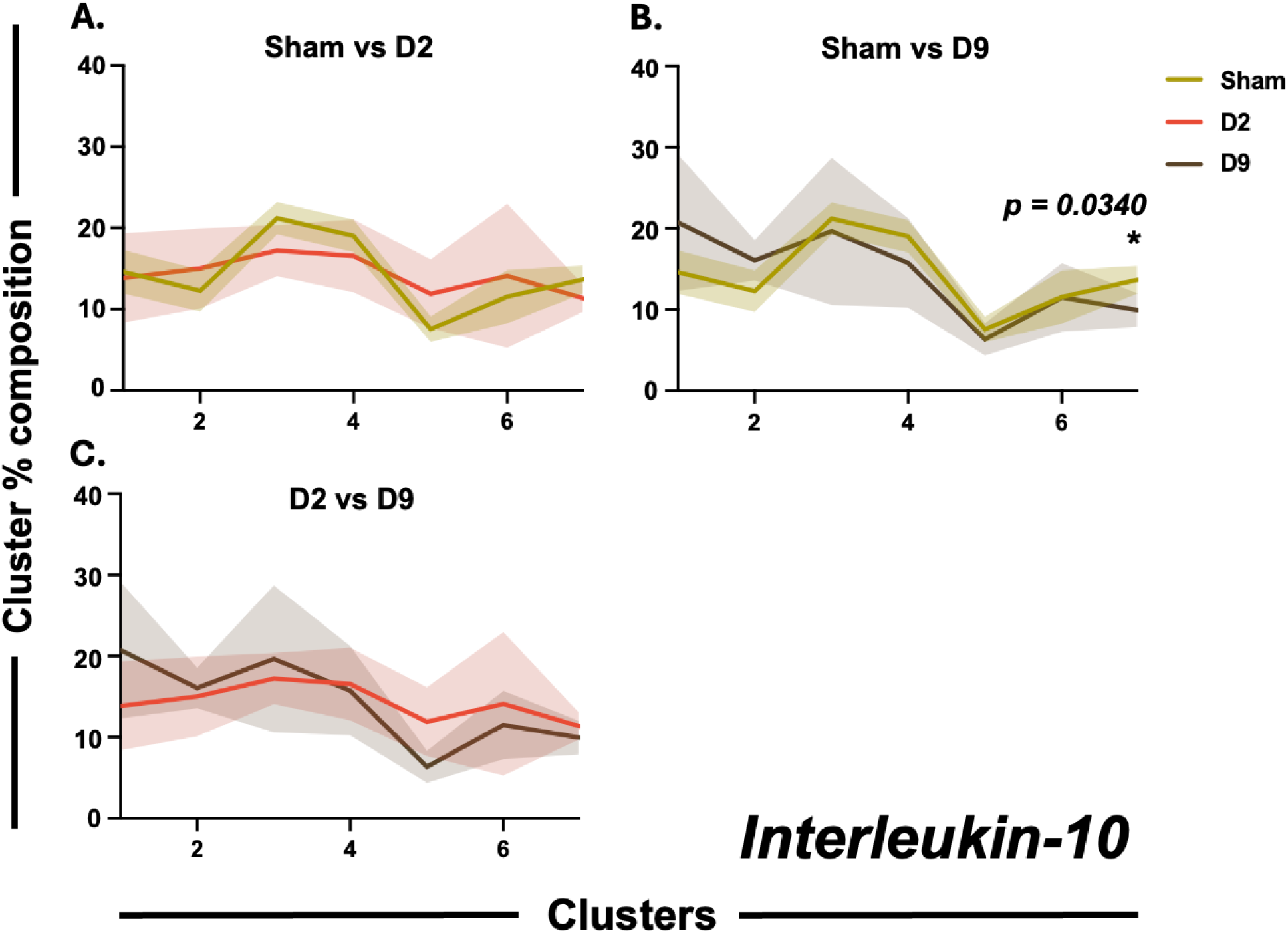
Proportion of reads in a given cluster based on condition is provided for the *Tgfb1* promoter. Flap-Enabled Next-Generation Capture (FENGC) of target sequences from F4/80^+^-enriched splenocytes at Day 2 (D2) and Day 9 (D9) was performed following 20% TBSA cutaneous burn injury in C57BL/6 mice. Standard deviation is provided by the shaded error bands.

**Supplemental Figure 3:**
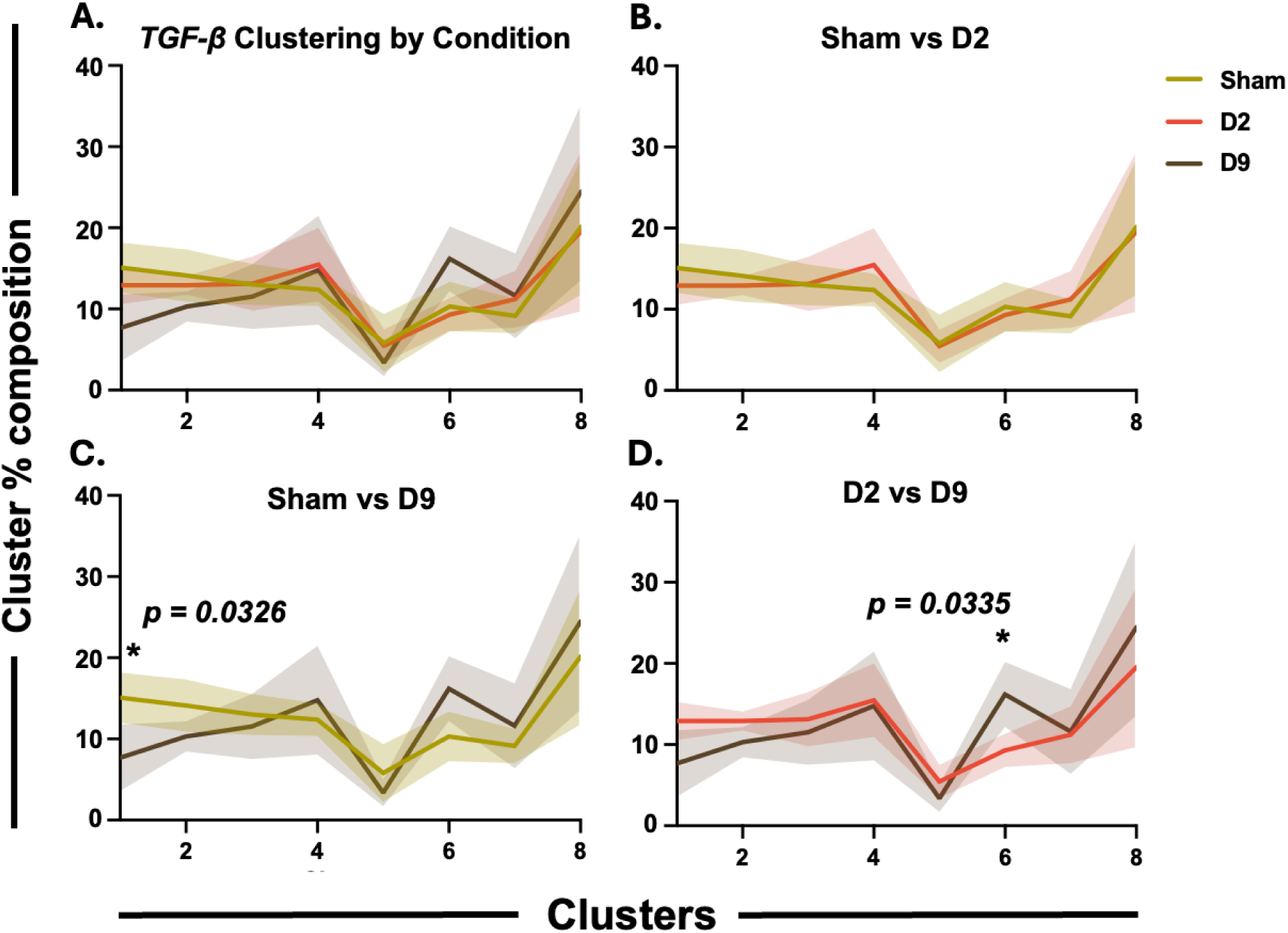
Proportion of reads in a given cluster based on condition is provided for the *Tgfb1* promoter. Flap-Enabled Next-Generation Capture (FENGC) of target sequences from F4/80^+^-enriched splenocytes at Day 2 (D2) and Day 9 (D9) was performed following 20% TBSA cutaneous burn injury in C57BL/6 mice. Standard deviation is provided by the shaded error bands.

**Supplemental Figure 4:**
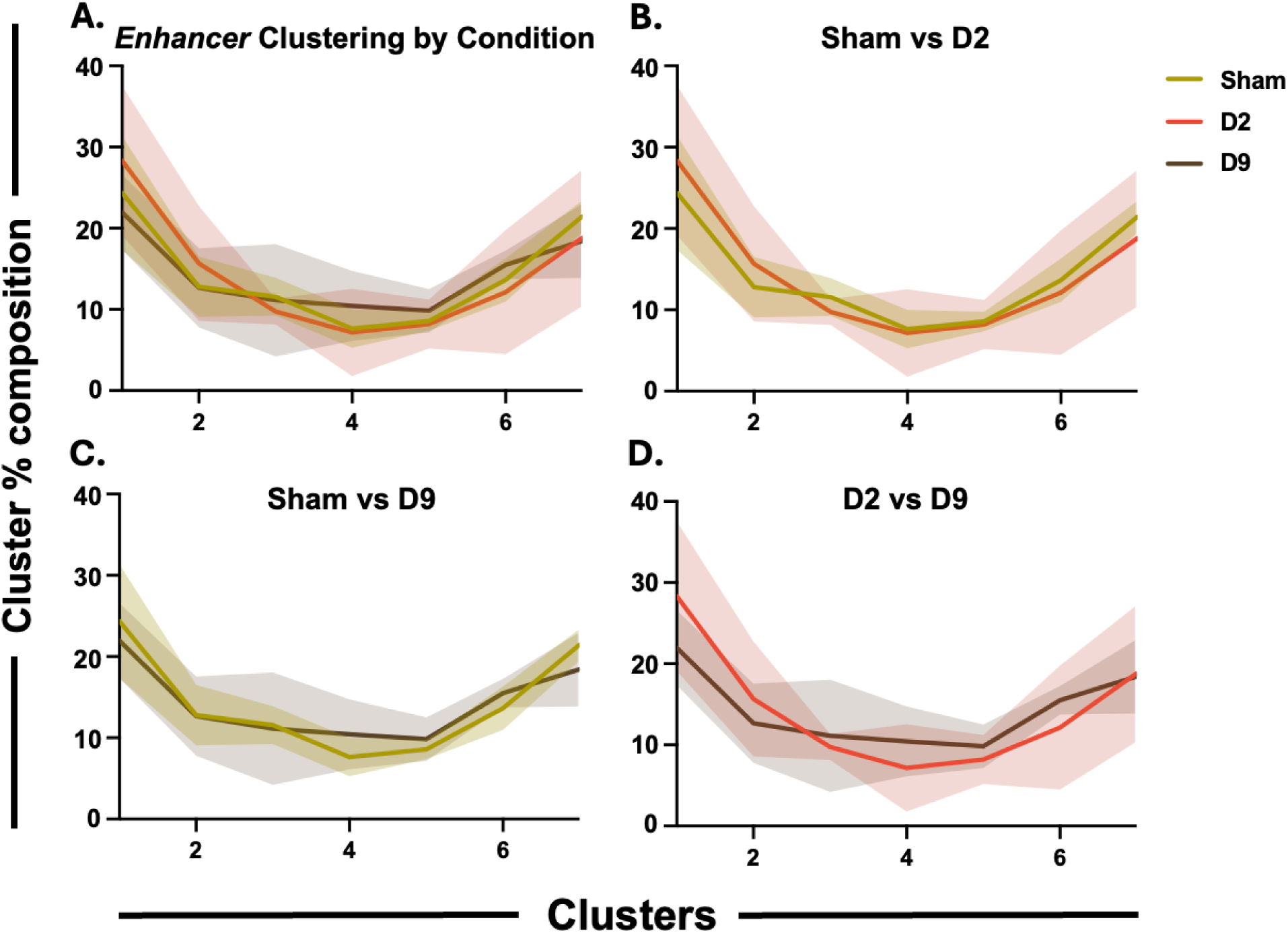
No difference in the total proportion of reads across Sham, D2 and D9 conditions. Flap-Enabled Next-Generation Capture (FENGC) of target sequences from F4/80^+^-enriched splenocytes at Day 2 (D2) and Day 9 (D9) was performed following 20% TBSA cutaneous burn injury in C57BL/6 mice. Standard deviation is provided by the shaded error bands.

**Supplemental Table 1:** Top 50 Upregulated and Downregulated Genes After Burn Injury. **Splenic tissue resident macrophages (trMø) harvested after burn injury exhibit specific immune and metabolic gene and pathway changes compared to trMø from sham injured mice:** mRNA isolated from splenic trMø 2, 9 or 14 days after 20% Total Body Surface Area burn injury, were tested for expression of immune and metabolic genes by nanoString analysis compared to trMø harvested from sham-injured mice. Data are presented as the log2-transformed differential fold change in immune or metabolic gene expression as shown in each label, with associated p-value significance (using Welch’s T test).

| Gene | Log2 Fold Change | -Log10(p-value) |
| --- | --- | --- |
| CCL5 | -5.2972 | 4.5460 |
| SLAMF7 | -5.2920 | 4.0164 |
| CD7 | -5.1699 | 3.6211 |
| KLRA8 | -5.0945 | 3.7663 |
| CCR6 | -5.0583 | 4.4965 |
| TCF7 | -4.8194 | 3.0319 |
| KLRD1 | -4.7562 | 3.7295 |
| CXCR5 | -4.6993 | 4.7616 |
| GZMA | -4.6989 | 5.4608 |
| THY1 | -4.6939 | 3.4062 |
| SPN | -4.6640 | 3.6484 |
| CXCR6 | -4.6630 | 3.8755 |
| CD3E | -4.5988 | 3.4857 |
| MAP4K1 | -4.4582 | 3.7229 |
| CD3D | -4.4544 | 3.6908 |
| LCK | -4.4374 | 3.0152 |
| GPR183 | -4.2673 | 4.1871 |
| CCR7 | -4.1744 | 4.2572 |
| CD40 | -4.1669 | 4.4017 |
| SH2D1A | -4.1588 | 12.3591 |
| CD8A | -4.0663 | 3.9296 |
| CCR2 | -4.0492 | 3.2266 |
| IL2RB | -4.0385 | 2.8393 |
| SIGIRR | -4.0000 | 2.8775 |
| CD27 | -3.9971 | 3.3699 |
| CR2 | -3.9441 | 3.4698 |
| TRAF1 | -3.9334 | 4.3472 |
| EOMES | -3.8745 | 2.8569 |
| IL7R | -3.8123 | 3.4283 |
| LTB | -3.7660 | 5.0657 |
| LEF1 | -3.7634 | 4.3735 |
| CD247 | -3.7283 | 4.4165 |
| IL1B | -3.6938 | 5.4189 |
| DPP4 | -3.6495 | 2.7211 |
| ITGAL | -3.6463 | 3.1717 |
| KLRA7 | -3.6301 | 3.0248 |
| KLRC1 | -3.6114 | 4.0715 |
| BTLA | -3.6080 | 4.7072 |
| IKBKE | -3.5255 | 4.1533 |
| CD19 | -3.4987 | 5.3405 |
| MS4A1 | -3.4946 | 5.3766 |
| CD226 | -3.4558 | 3.2383 |
| CXCR3 | -3.4497 | 2.9805 |
| ETS1 | -3.4364 | 5.3217 |
| CD69 | -3.4349 | 7.9923 |
| CD79A | -3.4150 | 2.8399 |
| TNFRSF13C | -3.4094 | 3.7155 |
| IL27RA | -3.3910 | 2.9786 |
| CD6 | -3.3716 | 10.9463 |
| XCL1 | -3.3542 | 4.2001 |
| LILRB4 | 1.6528 | 5.6116 |
| MARCO | 1.5656 | 1.9275 |
| IL18 | 1.4997 | 5.6611 |
| CMKLR1 | 1.3706 | 2.9994 |
| C1QB | 1.3436 | 5.7523 |
| CD163 | 1.3415 | 2.4586 |
| LTBR | 1.3410 | 2.2601 |
| VCAM1 | 1.2693 | 4.3502 |
| C1QA | 1.2638 | 5.2315 |
| PTK2 | 1.2630 | 1.5309 |
| PPARG | 1.2630 | 2.4112 |
| C6 | 1.2592 | 3.1792 |
| IL1A | 1.1635 | 2.1740 |
| TNFRSF11A | 1.1502 | 2.8087 |
| LAIR1 | 1.1407 | 4.8590 |

D2 Burn versus Sham - Metabolic Genes
| Gene | Log2 Fold Change | -Log10(p-value) |
| --- | --- | --- |
| LCK | -4.4896 | 2.2988 |
| NKG7 | -4.3796 | 3.2684 |
| KLRD1 | -4.3639 | 2.5304 |
| CD3E | -4.2788 | 1.9712 |
| BLK | -3.9764 | 2.8516 |
| CD3G | -3.9183 | 3.3819 |
| CTSW | -3.8366 | 1.9302 |
| CD3D | -3.8289 | 2.3754 |
| NCR1 | -3.6491 | 2.0912 |
| KMO | -3.6423 | 2.7809 |
| GZMA | -3.5850 | 3.3035 |
| CD27 | -3.5660 | 1.8736 |
| HSD11B1 | -3.4739 | 1.5166 |
| CCL5 | -3.4109 | 2.3815 |
| LAT | -3.3806 | 7.9855 |
| CD19 | -3.3777 | 3.5653 |
| CXCR6 | -3.3692 | 1.5166 |
| LTB | -3.3546 | 3.3035 |
| H2-OB | -3.3312 | 1.8736 |
| CD8A | -3.3219 | 1.8285 |
| MS4A1 | -3.2570 | 3.5653 |
| KYNU | -3.0959 | 3.7286 |
| CD28 | -3.0589 | 1.9678 |
| AKT3 | -2.8941 | 1.5069 |
| ITK | -2.8892 | 1.9712 |
| NDUFB11 | -2.4229 | 1.4324 |
| MYCL | -2.3062 | 1.5118 |
| ASS1 | -2.1775 | 1.4217 |
| HMOX1 | 2.1458 | 3.5063 |
| H2-EB1 | -2.1409 | 3.8410 |
| H2-AA | -2.0212 | 3.8410 |
| CXCL9 | -1.9987 | 2.3164 |
| CTSD | 1.9717 | 3.5306 |
| CTSL | 1.9391 | 1.8736 |
| IL2RA | -1.7801 | 3.8410 |
| FABP5 | 1.7435 | 1.4217 |
| CD163 | 1.7435 | 2.0765 |
| KMT2A | -1.6971 | 2.3164 |
| CD68 | 1.6802 | 1.4324 |
| APOE | 1.6531 | 3.2115 |
| ADA | 1.6001 | 1.4297 |
| EZH2 | -1.5996 | 1.4324 |
| CENPA | -1.5850 | 1.4864 |
| IDH1 | 1.5522 | 2.5967 |
| GLUL | 1.4850 | 3.3326 |
| CD247 | -1.4276 | 1.6281 |
| BIRC3 | -1.3312 | 1.3826 |
| PGD | 1.2893 | 2.1929 |
| HK3 | 1.2842 | 1.6495 |
| LTA4H | -1.2496 | 1.6098 |
| HEXA | 1.2172 | 1.6495 |
| H2-K1 | -1.1066 | 1.6962 |

D9 Burn versus Sham - Immune Genes
| Gene | Log2 Fold Change | -Log10(p-value) |
| --- | --- | --- |
| CTSG | 5.6682 | 3.4103 |
| CAMP | 4.6061 | 2.4589 |
| S100A9 | 3.6838 | 2.4589 |
| S100A8 | 3.4661 | 2.3493 |
| CLEC5A | 3.4150 | 2.0222 |

D9 Burn versus Sham - Metabolic Genes
| Gene | Log2 Fold Change | -Log10(p-value) |
| --- | --- | --- |
| MPO | 5.7101 | 6.1120 |
| PKLR | 3.0556 | 2.4312 |
| HDC | 2.9391 | 3.2779 |
| CPA3 | 2.6651 | 2.7488 |
| CDCA8 | 2.6455 | 3.2530 |
| RRM2 | 2.3770 | 2.1794 |
| ASNS | 2.3074 | 2.6249 |
| ALOX5 | 2.0496 | 3.0070 |
| CCNA2 | 1.9500 | 1.5140 |
| GDA | 1.9364 | 2.1551 |
| CCNB2 | 1.9279 | 1.8476 |
| MKI67 | 1.9180 | 2.4455 |
| COL4A2 | 1.8911 | 1.5484 |
| MGST3 | 1.8480 | 3.2469 |
| KIF2C | 1.8480 | 1.3802 |
| UBE2C | 1.8434 | 3.7287 |
| FOXM1 | 1.8349 | 1.9051 |
| PLK1 | 1.8342 | 1.6889 |
| FANCD2 | 1.8074 | 1.3806 |
| CLSPN | 1.8036 | 1.3524 |
| CD63 | 1.7757 | 3.9119 |
| UPP1 | 1.7655 | 1.6836 |
| BUB1 | 1.6781 | 1.8000 |
| BUB1B | 1.6702 | 1.8646 |
| GTSE1 | 1.6323 | 1.5372 |
| NCAPH | 1.6062 | 2.3757 |
| BRCA1 | 1.4854 | 1.8929 |
| PCLAF | 1.4531 | 2.2713 |
| ITGAM | 1.4318 | 1.5901 |
| EXO1 | 1.4099 | 1.4849 |
| DTL | 1.3857 | 1.9826 |
| TFRC | 1.3662 | 1.5213 |
| FANCA | 1.2630 | 1.4784 |
| CSF3R | 1.1570 | 1.4614 |
| TK1 | 1.1396 | 1.5985 |
| CD247 | -1.1307 | 1.4396 |

D14 burn versus Sham - Immune Genes
| Gene | Log2 Fold Change | -Log10(p-value) |
| --- | --- | --- |
| IFNA1 | 6.8298 | 14.6056 |
| CASP3 | -5.4811 | 4.3118 |
| SLAMF7 | -5.4514 | 3.9665 |
| MAP4K1 | -5.4460 | 4.2337 |
| CCL19 | 5.0417 | 6.8655 |
| RELA | -4.9900 | 4.1848 |
| CCL5 | -4.9753 | 4.2055 |
| NFATC1 | -4.7463 | 5.3354 |
| LEF1 | -4.6726 | 4.9692 |
| HIF1A | -4.6415 | 11.2315 |
| PSMC2 | -4.6293 | 3.2530 |
| IL28A | 4.5699 | 3.2629 |
| C1S | 4.5383 | 6.5236 |
| IL25 | 4.5220 | 4.8045 |
| FN1 | -4.4739 | 2.9002 |
| CCL2 | 4.4429 | 3.0429 |
| IL9 | 4.4429 | 3.1873 |
| TRAF1 | -4.4258 | 4.5979 |
| CCL11 | 4.3038 | 2.4845 |
| CD109 | 4.2752 | 3.7708 |
| IKZF4 | 4.1997 | 3.4658 |
| SH2D1A | -4.1588 | 12.1843 |
| KIR3DL2 | 4.0461 | 5.4436 |
| CXCR5 | -3.9413 | 4.2321 |
| IL21R | -3.8628 | 4.1092 |
| MX1 | 3.8493 | 7.0629 |
| LCK | -3.8379 | 2.5152 |
| TNFSF15 | 3.7814 | 2.7049 |
| THY1 | -3.7590 | 2.7282 |
| AHR | -3.7370 | 2.5735 |
| IFI35 | -3.7302 | 4.1900 |
| H2-AB1 | -3.6829 | 6.8868 |
| TRAF6 | -3.6818 | 3.2110 |
| FCER1A | 3.6370 | 6.1657 |
| ABL1 | -3.6235 | 4.5200 |
| CX3CR1 | -3.5858 | 3.9811 |
| CCL20 | 3.5850 | 1.8143 |
| ICAM4 | 3.5850 | 2.0742 |
| CD69 | -3.5742 | 7.9729 |
| LTF | 3.5622 | 2.6331 |
| C8B | 3.5547 | 5.8031 |
| H2-EB1 | -3.5518 | 5.2724 |
| BAX | -3.5005 | 4.6932 |
| CD74 | -3.4892 | 7.2895 |
| CCR10 | 3.4215 | 2.4795 |
| NCF4 | -3.4150 | 2.9888 |
| NCAM1 | 3.4147 | 6.9619 |
| IL2RB | -3.4133 | 2.2759 |
| IKZF3 | -3.3891 | 4.3066 |
| LTB | -3.3785 | 4.5509 |
| CD6 | -3.3716 | 10.7720 |
| SERPING1 | 3.3576 | 2.3200 |
| TCF7 | -3.3370 | 2.0518 |
| CCR7 | -3.3331 | 3.5053 |
| IL12B | 3.3219 | 2.4251 |
| ITGA4 | -3.3116 | 4.1444 |
| ITGAM | -3.2594 | 3.6892 |
| C7 | 3.2574 | 1.7994 |
| IL1B | -3.2528 | 4.8070 |
| IL23R | 3.2479 | 2.2344 |
| DEFB14 | 3.2479 | 2.1137 |
| IL6ST | -3.2224 | 2.4781 |
| TGFB3 | 3.2185 | 1.9911 |
| ITGAL | -3.2175 | 2.7090 |
| BTNL1 | 3.2057 | 2.0346 |
| PLA2G2E | 3.1699 | 1.8158 |
| IL2 | 3.1610 | 1.9543 |
| IL6 | 3.1601 | 2.1029 |
| ARHGDIB | -3.1543 | 8.3398 |
| FCAMR | 3.1396 | 2.0116 |
| TAPBP | -3.1155 | 4.7824 |
| H2-AA | -3.1032 | 6.4469 |
| TNFRSF13C | -3.0875 | 3.3957 |
| CD3D | -3.0780 | 2.5946 |
| CYBB | -3.0366 | 5.2779 |
| TRAF2 | -3.0080 | 2.9919 |
| FYN | -3.0069 | 5.9059 |
| TSLP | 2.9600 | 2.2674 |
| BCL3 | -2.9271 | 3.3964 |
| IL1R2 | 2.9069 | 1.5433 |
| TNFRSF11A | 2.8858 | 5.7005 |
| STAT2 | -2.8704 | 3.5264 |
| MUC1 | 2.8697 | 1.9728 |
| SELL | -2.8577 | 2.0042 |
| GZMA | -2.8534 | 4.1184 |
| CFH | 2.8479 | 3.7455 |
| MASP2 | 2.7692 | 2.0741 |
| IL22RA2 | 2.7549 | 1.7624 |
| IL21 | 2.7257 | 2.0796 |
| TNFRSF17 | 2.7199 | 1.8428 |
| CD59B | 2.6965 | 2.0617 |
| DEFB1 | 2.6384 | 2.0731 |
| NOS2 | 2.5480 | 1.5450 |
| IL1RL1 | 2.5460 | 2.7867 |
| SRC | 2.5349 | 2.6313 |
| IFITM1 | 2.3854 | 2.9992 |
| TRAF4 | 2.3252 | 4.2662 |
| IRF3 | 2.3219 | 1.5411 |
| CD28 | 2.3060 | 2.1667 |
| CTSG | 2.2288 | 1.4348 |

**Supplemental Table 1: Splenic tissue resident macrophages (trMø) harvested after burn injury exhibit specific immune and metabolic gene and pathway changes compared to trMø from sham injured mice:** mRNA isolated from splenic trMø 2, 9 or 14 days after 20% Total Body Surface Area burn injury, were tested for expression of immune and metabolic genes by nanoString analysis compared to trMø harvested from sham-injured mice.
| Gene | Log2 Fold Change | -Log10(p-value) |
| --- | --- | --- |
| TYMS | 5.5714 | 3.0992 |
| NKG7 | -5.2116 | 3.6729 |
| PSMA7 | -4.7004 | 2.8828 |
| PDP1 | 4.6348 | 3.1958 |
| LCK | -4.5688 | 2.5082 |
| G6PC | 4.5392 | 2.2054 |
| COX4I1 | -4.4594 | 3.1958 |
| ZFP975 | 4.2216 | 3.0992 |
| PTGES | 4.1898 | 1.6431 |
| GZMA | -4.1699 | 3.1254 |
| BHMT | 4.1699 | 1.6370 |
| CDC20 | 4.0959 | 2.5607 |
| NOS2 | 4.0928 | 2.1804 |
| TPH1 | 4.0661 | 1.5257 |
| STAT5B | -4.0027 | 3.4314 |
| CD209E | 4.0000 | 1.9535 |
| VPS28 | -3.9151 | 2.6597 |
| SLC16A8 | 3.8826 | 1.9613 |
| IMPDH2 | -3.8594 | 2.5607 |
| CCL19 | 3.8580 | 2.0389 |
| COL6A2 | 3.8580 | 1.9389 |
| PDPK1 | -3.7886 | 4.0717 |
| KIF2C | 3.7142 | 2.0295 |
| RRM1 | -3.6630 | 2.6458 |
| TYMP | 3.6511 | 1.8427 |
| HIF3A | 3.6439 | 1.4758 |
| TCL1 | 3.6439 | 1.4962 |
| ITK | -3.6355 | 2.6342 |
| MYBL2 | 3.5869 | 2.5727 |
| KLRB1A | 3.5850 | 1.9613 |
| HJURP | 3.5699 | 2.4857 |
| EFNA4 | 3.5546 | 1.4997 |
| BCL2L1 | 3.5258 | 3.1254 |
| AKR1C6 | 3.5236 | 1.5209 |
| PDK1 | -3.5197 | 2.0882 |
| VEGFA | 3.4919 | 2.0389 |
| SERINC2 | 3.4919 | 1.5279 |
| DMGDH | 3.4594 | 1.3585 |
| LAMTOR4 | 3.4594 | 1.4298 |
| LAT | -3.4531 | 7.8938 |
| H2-AA | -3.4146 | 5.3778 |
| HK2 | 3.3947 | 2.2130 |
| CACNG3 | 3.3923 | 1.3702 |
| CCL5 | -3.3785 | 2.5607 |
| NEDD8 | -3.3614 | 2.3112 |
| SLC6A12 | 3.3576 | 1.4298 |
| SCD1 | -3.3536 | 2.6180 |
| GZMB | 3.3505 | 2.0834 |
| IL2 | 3.3219 | 1.4298 |
| DDC | 3.3219 | 1.5257 |
| PHGDH | 3.2956 | 2.5607 |
| TRAF1 | -3.2912 | 3.0992 |
| ACMSD | 3.2854 | 1.5257 |
| NAALAD2 | 3.2479 | 1.6741 |
| DCK | -3.2301 | 3.5626 |
| H2-EB1 | -3.2119 | 5.1111 |
| SIGLECF | 3.2095 | 1.5346 |
| H2-D1 | -3.2052 | 2.7031 |
| CYBB | -3.1710 | 3.8434 |
| SREBF2 | -3.1570 | 2.7476 |
| CXCL9 | -3.1521 | 3.4654 |
| SERINC3 | -3.1102 | 3.7555 |
| RPS6KA1 | -3.1031 | 2.9756 |
| CENPA | -3.0995 | 2.2587 |
| FCER1A | 3.0875 | 1.3665 |
| CYP1B1 | 3.0875 | 1.3665 |
| CYP8B1 | 3.0815 | 1.8494 |
| INMT | 3.0444 | 1.4754 |
| KYAT1 | 3.0444 | 1.9553 |
| IFNG | 3.0112 | 1.3185 |
| DAO | 3.0000 | 1.3090 |
| WNT1 | 3.0000 | 1.3665 |
| NOS3 | 3.0000 | 1.3799 |
| LTA4H | -2.9866 | 2.9756 |
| TPPA | 2.9542 | 1.3879 |
| TDO2 | 2.9542 | 1.3185 |
| SLC2A5 | 2.9542 | 1.3432 |
| IDO2 | 2.9367 | 2.0843 |
| PSMD13 | -2.9366 | 2.4071 |
| LTB | -2.9117 | 3.0539 |
| KYNU | -2.8888 | 3.7555 |
| OAT | -2.8248 | 1.7079 |
| CD27 | -2.8019 | 1.6359 |
| JAK2 | -2.7997 | 1.6925 |
| TRAF6 | -2.7726 | 1.9286 |
| RPLP0 | -2.6696 | 5.3778 |
| EPC1 | -2.6660 | 1.3973 |
| NFKB1 | -2.6630 | 2.3289 |
| CD180 | -2.6541 | 2.8652 |
| NRAS | -2.6522 | 1.7513 |
| CD3G | -2.5964 | 3.2177 |
| GOT1 | -2.5922 | 2.4909 |
| NDUFB10 | -2.5703 | 3.0665 |
| GART | -2.5558 | 1.9877 |
| PTPRC | -2.5332 | 3.7555 |
| NCR1 | -2.5271 | 1.5915 |
| HEXA | -2.5219 | 2.7240 |
| CTSS | -2.5192 | 3.2177 |
| H2-K1 | -2.5011 | 3.5626 |
| CTSW | -2.4914 | 1.3849 |

## Supplemental Materials 1

### Methyltransferase Accessibility Protocol for Individual Templates combined with Flap-Enabled Next-Generation Capture (MAPit-FENGC) epigenetic analysis

One million cells purified from each mouse were washed with ice-cold PBS, pH 7.2. Cells were centrifuged at 1,000 x g for 5 min, then washed in ice-cold cell resuspension buffer (20 mM HEPES pH 7.5, 70 mM NaCl, 0.25 mM EDTA pH 8.0, 0.5 mM EGTA pH 8.0, 0.5% glycerol, freshly supplemented with 10 mM DTT and 0.25 mM PMSF). Cells were centrifuged at 1,000 x g for 5 min before being resuspended with 92 μL of resuspension buffer with 0.5 % digitonin. Cells were then stained with trypan blue to ascertain 100 % permeabilization. The cells were then treated with 100 U M.CviPI GpC methyltransferase (100 U/million cells; New England Biolabs, M0227B-HI) with fresh 160 µM S-adenosyl-L-methionine for 15 min at 37°C. The reaction was terminated by adding an equal volume of 10 mM EDTA, 100 mM NaCl, and 1% (w/v) SDS followed by a quick vortex at medium speed. The nuclei were treated with 100 µg/ml RNase A for 30 min at 37°C followed with 100 µg/ml proteinase K treatment overnight at 50°C. Extraction of genomic DNA was conducted using phenol-chloroform-isoamyl alcohol (25:24:1, v/v) phase separation, which was followed by ethanol precipitation, then resuspension in molecular-grade H_2_O. The full method is available at https://www.biorxiv.org/content/10.1101/2022.11.08.515732v1.full.

